# Disrupted Developmental Trajectory of Ultrasonic Vocalizations in a Rat Model of Fragile X Syndrome

**DOI:** 10.64898/2026.07.31.742126

**Authors:** D. Walker Gauthier, Arjun Vaidya, Benjamin D. Auerbach

## Abstract

Communication deficits are a defining feature of autism spectrum disorder (ASD) and among the earliest detectable markers of atypical neurodevelopment. Yet how specific genetic ASD risk factors shape the developmental trajectory of vocal communication remains poorly understood. Fragile X syndrome (FXS) is the most common inherited cause of ASD, resulting from the transcriptional silencing of the FMR1 gene, and a majority of FXS individuals exhibit impaired language development and atypical vocal communication. Rodent ultrasonic vocalizations (USVs) produced during maternal isolation provide a tractable model for studying the developmental trajectory of early vocal communication in FXS. Here, we characterized isolation- induced USVs in *Fmr1* knockout (KO) and littermate wildtype (WT) rats from postnatal days 3– 21 to determine whether *Fmr1* mutation disrupts the acoustic structure, temporal organization, or sequential syntax of USVs across postnatal development. We found that *Fmr1* KO rat pups exhibited reduced call number during the peak developmental window for isolation-induced calling (p6–p10), while acoustic structure and temporal organization were largely preserved. Network analysis of call transitions revealed that WT pups exhibited a progressive increase in syntactic complexity from p3–p10. However, this developmental trajectory was significantly altered and delayed in *Fmr1* KO pups. Together, these findings demonstrate that *Fmr1* mutation not only disrupts vocal production but the developmental expansion of syntactic flexibility in rats, highlighting USV syntax as a sensitive marker of atypical communicative development in FXS models.

## INTRODUCTION

Autism spectrum disorders (ASD) are a set of neurodevelopmental conditions defined by persistent deficits in social communication and restricted, repetitive patterns of behavior^1^. Vocal communication impairments are among the most common and clinically significant features of ASD, encompassing delayed or absent speech, echolalia, and atypical prosody^2–4^. These communication impairments are also dynamic, emerging early in postnatal development and following atypical trajectories, making them among the strongest predictors of long-term functional outcome for ASD individuals^5,6^. Understanding how genetic ASD risk factors give rise to these atypical trajectories therefore requires model systems capable of capturing the acoustic structure and temporal organization of vocal output across the early postnatal period.

Rodent ultrasonic vocalizations (USVs) provide a tractable, non-invasive window into the development of affective communication^7,8^. In the isolation paradigm, pups separated from the dam emit USVs as a distress signal, eliciting maternal search-and-retrieval behavior in a manner functionally analogous to infant crying^9,10^. The calls produced by rat pups are acoustically heterogeneous, comprising distinct subtypes with characteristic temporal and spectral features that may carry differential communicative value^11,12^. Critically, USV production in rat pups follows a well-characterized developmental trajectory across the first three postnatal weeks, with systematic changes in call number, acoustic properties, and call type composition^13,14^. This developmental arc has made the isolation USV paradigm a widely adopted tool for characterizing communicative phenotypes in rodent models of ASD^11,15^.

Beyond individual call properties, the sequential organization of USV output constitutes an additional and largely independent dimension of vocal communication^15,16^. Markov chain-based transition analyses and graph-theoretic approaches have revealed that USV sequences contain statistically structured patterns that influence maternal behavior, with more predictable and temporally coherent sequences eliciting stronger retrieval responses than random or degraded sequences^16–18^. Syntactic complexity is particularly relevant to ASD, in which restricted inflexible behavioral repertoires are a defining feature^19,20^. Despite its potential utility for examining communication deficits, the developmental trajectory of USV syntactic complexity has received comparatively little attention in genetic rodent models of ASD.

Fragile X syndrome (FXS) is the most common inherited cause of ASD and intellectual disability, arising from the transcriptional silencing of FMR1 gene and consequent loss of its protein product, fragile X messenger ribonucleoprotein (FMRP)^21,22^. Like the greater autistic population, FXS individuals often exhibit significant language delays, with deficits in expressive communication emerging as early as the first year of life, including reduced babbling and atypical early vocalization^23,24^. Importantly, the defined genetic etiology and well-characterized molecular pathology of FXS have enabled the development of highly tractable rodent models, including the *Fmr1* knockout (KO) rat, that recapitulate core features of the syndrome^25^, including abnormalities in USV production^26^. However, the temporal organization and syntactic structure of vocal output in Fmr1 KO animals remains largely uncharacterized, with most studies focusing on quantitative measures such as call number and acoustic properties, often at a single developmental time-point^27–29^. Whether loss of FMRP disrupts the sequential architecture of early vocal communication— and how any such disruption unfolds across postnatal development — remains unclear.

Here we characterized the developmental trajectory of isolation-induced USVs in littermate *Fmr1* KO and wildtype (WT) rats from postnatal day 3 to 21, examining acoustic properties, temporal patterning, call type composition, and syntactic complexity. We found a transient reduction in total call number in Fmr1 KO rats during the peak developmental window for isolation-induced calling (p6–p10), while the acoustic properties and temporal organization of calls remained largely preserved across all developmental timepoints. Strikingly, whereas WT animals exhibited a progressive increase in syntactic complexity across the first three postnatal weeks, this developmental trajectory was markedly attenuated in *Fmr1* KO pups. These findings indicate that loss of FMRP disrupts the organizational structure of early vocal output despite largely preserved acoustic properties, pointing to syntactic complexity as a potential metric for detecting communicative disruption in rodent models of ASD.

## RESULTS

### Acoustic properties of isolation-induced USVs are unaltered in *Fmr1* KO rats

Atypical acoustic properties of vocal output have been documented in individuals with ASD, including differences in fundamental frequency, pitch variability, and speech intensity relative to neurotypical controls^30,31^. Notably, these acoustic differences are detectable as early as infancy, with at-risk infants later diagnosed with ASD producing cries with higher and more variable fundamental frequency than low-risk controls^32^. To investigate whether analogous acoustic differences in distress vocalizations are present in a rat model of FXS across postnatal development, *Fmr1* KO and WT pups were isolated from the dam at postnatal days (p) 3, 6, 10, 14, and 21. Isolation-induced USVs were recorded using an ultrasonic microphone, with call detection, feature extraction, and classification performed with DeepSqueak^33^ (Fig 1).

**Figure 1.**
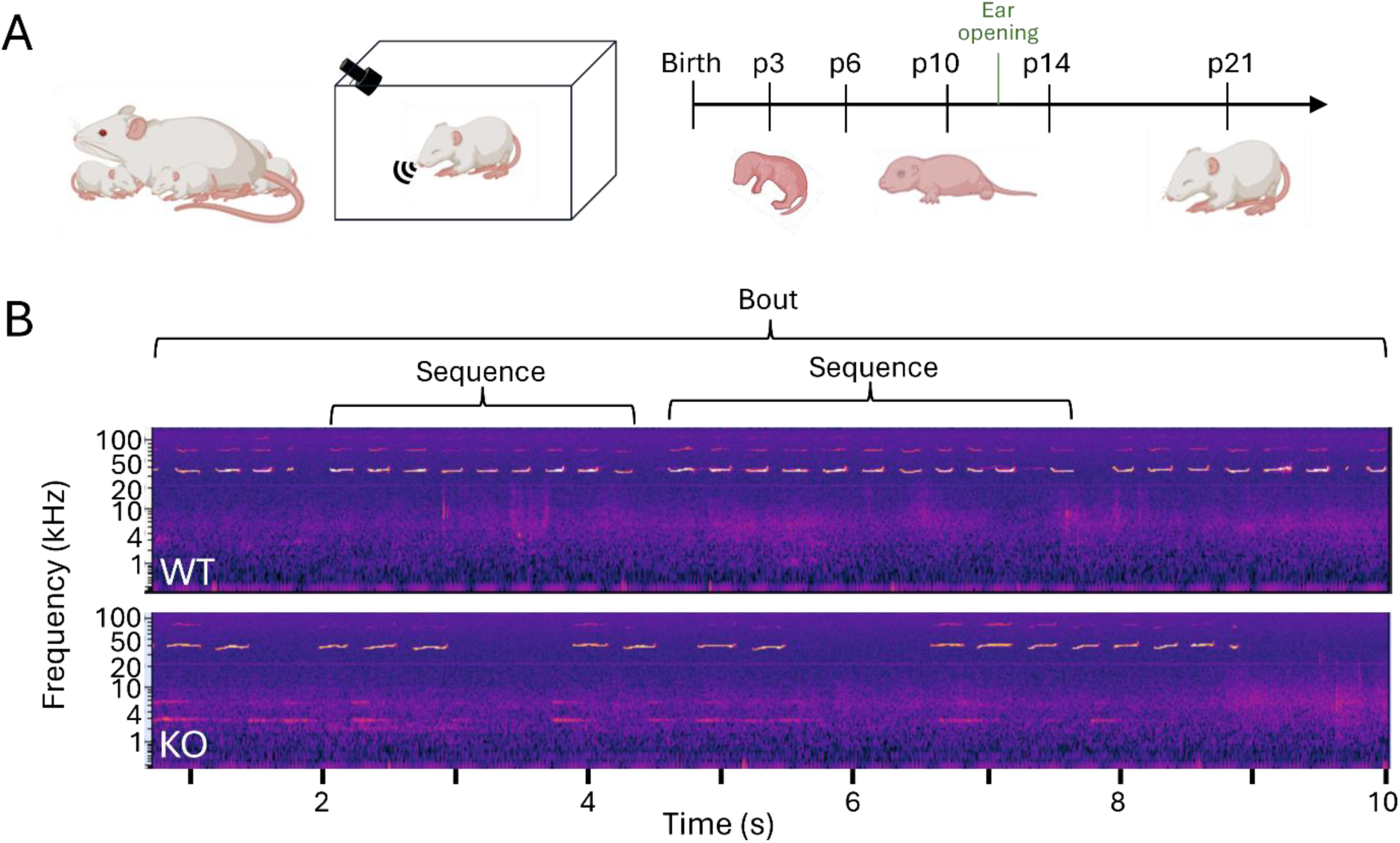
Maternal isolation-induced USV recording paradigm. **(A)** Schematic of the maternal isolation paradigm. Individual rat pups were removed from the home cage and dam and placed in a sound-attenuating recording chamber for a 5-minute isolation session. Ultrasonic vocalizations were recorded using an ultrasonic microphone and detected and analyzed using DeepSqueak. Recordings were conducted at postnatal days 3, 6, 10, 14, and 21. **(B)** USV calls from an example WT (top) and Fmr1 KO (bottom) pup at post- natal day 6 (p6).

We quantified the acoustic properties of USVs produced by each genotype across developmental timepoints, including call intensity (measured as mean power), call length, principal frequency, and bandwidth (measured as delta frequency) (Fig 2). Physical call properties varied significantly across development, with call intensity, duration, and bandwidth steadily increasing from p3 to p14 while principal frequency initially decreased from p3 to p10 then increased from p14 onward. These trends are consistent with previously characterized vocal development trajectory in rat pups^34^. However, no genotype differences were observed in call intensity (Fig. 2A; Two-Way ANOVA (genotype): F = 1.827, p = 0.3092), call length (Fig. 2B; Scheirer-Ray-Hare (genotype): H = 0.5787, p =0.4468), principal frequency (Fig. 2C; Scheirer-Ray-Hare (genotype): H = 0.0593, p = 0.8077), or bandwidth (Fig. 2D; Scheirer-Ray-Hare (genotype): H = 2.4083, p = 0.1207) at any developmental timepoint. Taken together, these results indicate that the basic acoustic structure of isolation-induced USVs develops normally in *Fmr1* KO rats, suggesting that loss of FMRP does not disrupt the fundamental motor or laryngeal mechanisms underlying call production during the early postnatal period.

**Figure 2:**
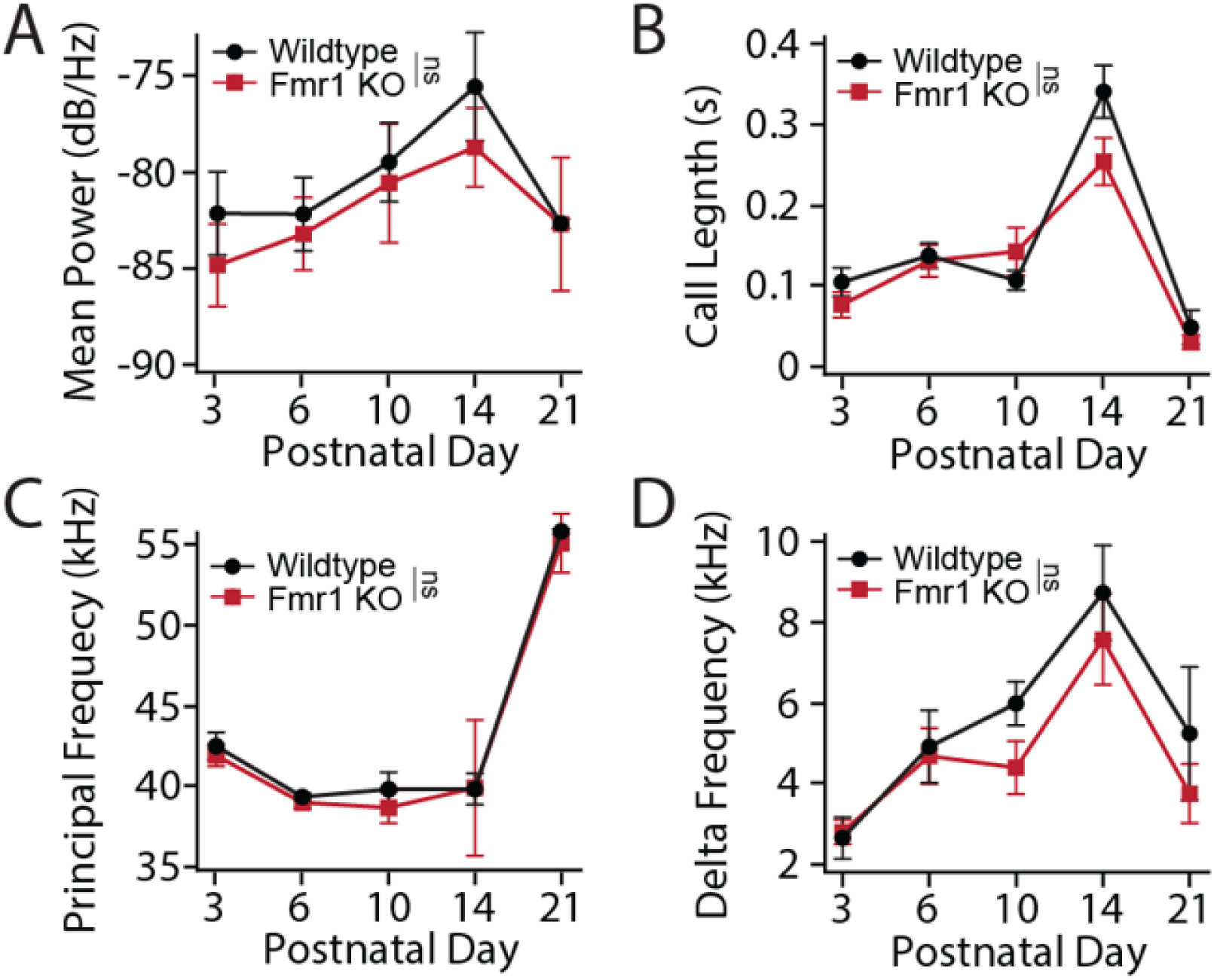
Physical properties of isolation-induced USVs are preserved across development in Fmr1 KO rats. **(A)** Mean call power (dB/Hz) across postnatal development in WT (black) and Fmr1 KO (red) rats. **(B)** Call length (s) across postnatal development. **(C)** Principal frequency (kHz) across postnatal development. **(D)** Delta frequency (kHz), reflecting the spectral bandwidth of each call, across postnatal development. All values are means ± SEM. ns = not significant.

### *Fmr1* KO pups produce fewer isolation-induced USVs but maintain temporal calling structure

Reduced frequency of communicative vocalizations is among the earliest detectable markers of ASD risk, with infants later diagnosed with ASD producing fewer speech-related vocalizations in the first two years of life relative to typically developing peers^35,36^. Moreover, FXS infants have been shown to exhibit less syllables and reduced canonical babbling ratios compared to typically developing infants^37^. We therefore quantified total call number in WT and *Fmr1* KO rat pups across postnatal age (Fig 3A-C). Total call number followed a characteristic developmental profile, with an inverted U pattern that peaks around p10. We found that *Fmr1* KO rat pups produced significantly fewer calls than WT animals across this peak calling period (Fig. 3A; Scheirer-Ray-Hare (genotype): H = 5.5526, *p = 0.0187), with a particularly pronounced decrease at p6. This reduction persisted across higher order temporal groupings (Fig 1B), with reduced number of call sequences in *Fmr1* KO pups as well (Fig 3B; Scheirer-Ray-Hare (genotype): H = 4.2048, *p = 0.0403). A similar trend was observed for bout number, though this did not reach statistical significance (Fig 3C; Scheirer-Ray-Hare (genotype), H = 2.9967, p = 0.0834). This reduction in vocal output is consistent with FXS infant data^37,38^, indicating that altered isolation-induced USV call rate is an evolutionarily conserved and translationally relevant FXS phenotype.

**Figure 3:**
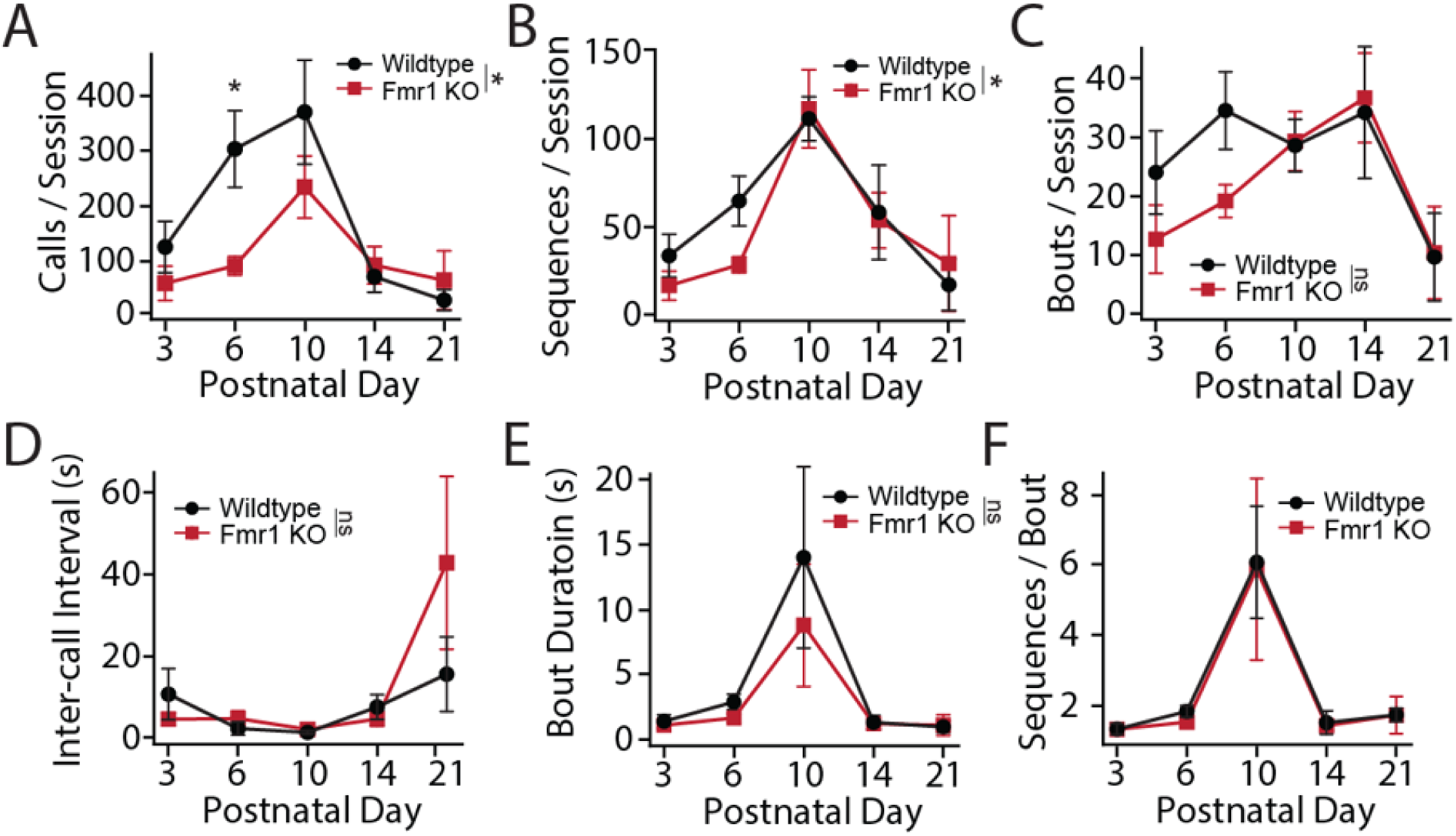
Fmr1 KO pups produce fewer isolation calls but maintain temporal calling structure. **(A)** Total call number per 5-minute isolation session across postnatal development in WT (black) and Fmr1 KO (red) rats. **(B)** Total sequence number per 5-minute isolation session across postnatal development in WT (black) and Fmr1 KO (red) rats. **(C)** Total number of bouts per 5- minute isolation session across postnatal development in WT (black) and Fmr1 KO (red) rats. **(D)** Mean inter-call interval (ICI; seconds) across postnatal development. **(E)** Mean bout duration (seconds) across postnatal development. **(F)** Mean number of sequences per bout across postnatal development. All values are means ± SEM. *p < 0.05, ns = not significant.

We next assessed the temporal organization of USV calls in WT and *Fmr1* pups (Fig 1B). USV temporal structure significantly varied by postnatal age, with bout duration (Scheirer-Ray-Hare (PND): H = 19.577, **p = 0.001) and sequences per bout (Scheirer-Ray-Hare (PND): H = 27.636, ***p < 0.0001)) following a similar inverted U pattern to that observed for overall call number. However, we did not observe any clear genotype differences in higher order temporal structure of pup calls at any age. Inter-call interval did not significantly differ between genotypes at any developmental timepoint (Fig. 3D; Scheirer-Ray-Hare (genotype): H = 1.6141, p = 0.2039). Bout duration (Fig. 3E; Scheirer-Ray-Hare (genotype): H = 2.7562, p = 0.0969) and the number of sequences per bout (Fig. 3F; Scheirer-Ray-Hare (genotype):H = 2.532, p = 0.1116) were similarly unaltered. Together, these results indicate that the higher-order temporal clustering of calls into bouts and sequences develops on a normal trajectory in *Fmr1* KO rats, despite alterations in total vocal output.

### *Fmr1* KO pups show attenuated development of syntactic complexity

Beyond the acoustic and temporal properties of individual calls, the sequential structure of USV output represents an additional dimension of communicative complexity that may be particularly sensitive to neurodevelopmental disruption^17,39^. To examine call complexity, we performed supervised clustering of call type using DeepSqueak region based convolutional neural network^33^ (Fig 4A). The predominant call type observed at early developmental ages were flat calls (Fig 4B), which are thought to be the rodent pup homolog of aversive distress vocalizations^40,41^. There was a reduction in the relative proportion of flat calls across development in both genotypes, consistent with prior reports of decreased stress-related vocal output as pups mature and become less dependent on maternal care^14^. This reduction coincided with an increase in the number of short and spectrally complex calls, which are thought to be associated with prosocial behaviors^13^. *Fmr1* KO rat pups exhibited a relative delay in the reduction of flat calls relative to short and other complex calls (Fig. 4A; Chi2 test; p3: chi2 = 5.065, p = 0.08; p6: chi2 = 15.901, ***p = 0.0003; p10: chi2 = 142.567, ***p < 0.0001). This result indicates that *Fmr1* KO pups show delayed maturation of call type composition, raising the question of whether the sequential structure of vocal output is similarly disrupted.

**Figure 4:**
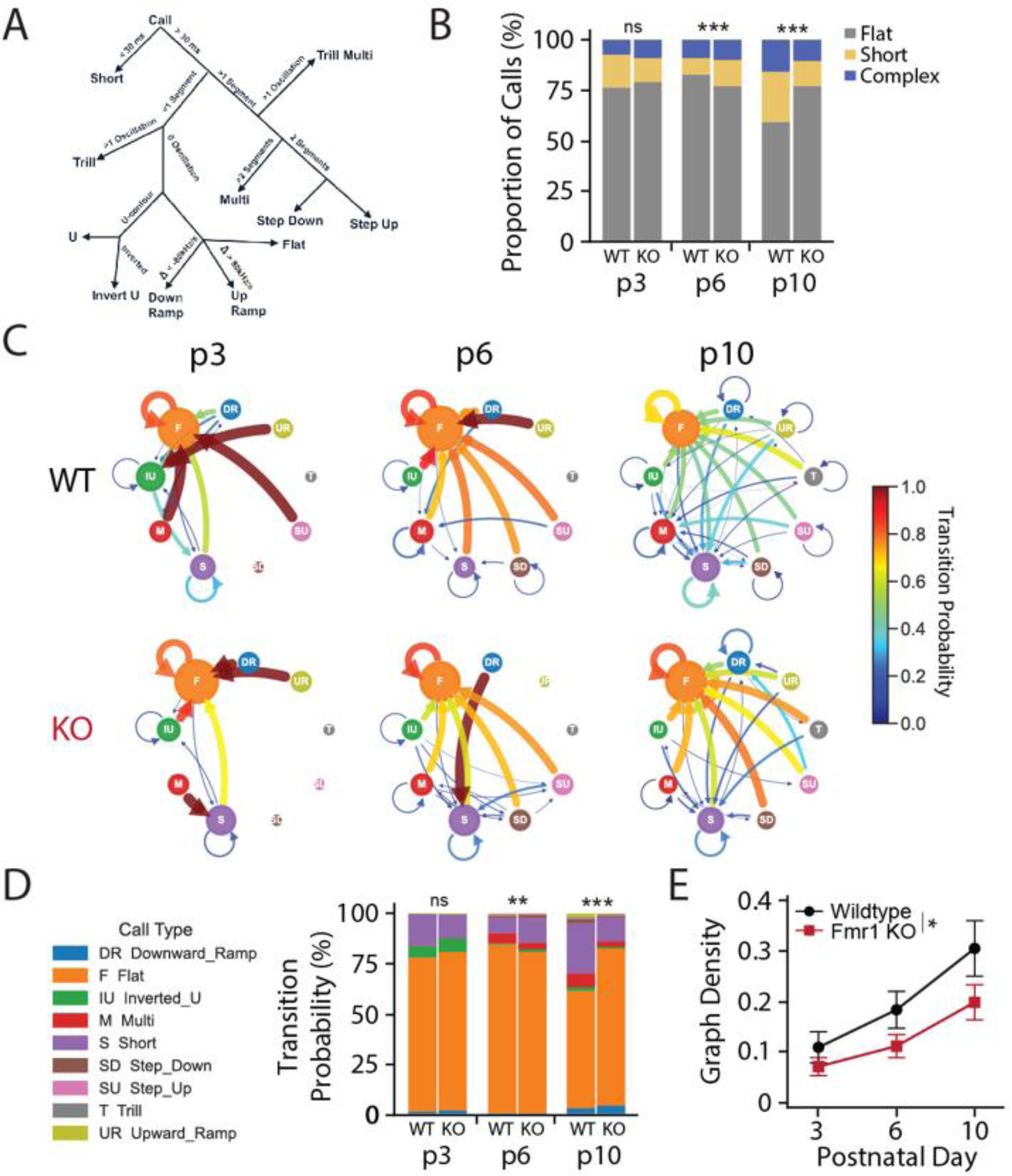
Fmr1 KO pups show an attenuated development of syntactic complexity. **(A)** Decision tree used for call labeling for region based convolutional neural network training. **(B)** Stacked bar plots showing the proportions of 3 broad call type categorizations (flat, short, complex) for WT (left bar) and Fmr1 KO (right bar) pups at postnatal days 3, 6, and 10. Call types are rank-ordered within each bar from most to least frequent. **(C)** Call type transition probability networks for wildtype (top row) and Fmr1 KO (bottom row) pups at each developmental timepoint. Node size reflects the relative frequency of each call type; edge width and color reflects the transition probability between call types. Only transitions edges with probability > 0.03 are shown. **(D)** Incoming transition probability stacked bar graph showing the marginal probability of transitioning to each call type for each genotype for each postnatal day. **(E)** Transition network graph density across postnatal development in wildtype (black) and Fmr1 KO (red) rats. Data from p14 and p21 are shown for completeness but were excluded from statistical analysis due to reduced call numbers at these timepoints. All values for line plots represent means ± SEM. *p < 0.05, **p < 0.01, ***p < 0.001, ns = not significant.

To examine whether genotype differences existed at the level of sequencing, calls were further classified into 11 call types based on their physical characteristics (Fig 4A) and transition probability networks were constructed for each genotype at each developmental timepoint (Fig. 4C). This graph network over all call types represents the probability of transitioning between each pair of call types within a bout. We restricted our analysis to data from p3 to p10, as call number was substantially reduced following ear opening (Fig 3A), precluding reliable sequence analysis. Visual inspection of these networks revealed qualitative differences in the richness and directionality of transitions between genotypes, motivating a quantitative analysis of transition network structure. First, the summed total conditional probability for each call was compared between genotypes (Fig 4D). There was a significant difference in these distributions at both p6 and p10 (Chi2 test; p3: chi2 = 3.851, p = 0.5710; p6: chi2 = 21.15, **p = 0.0053; p10: chi2 = 184.527, ***p < 0.0001). At early timepoints (p3–p6), transitions were biased toward flat calls in both genotypes. By p10, this call transition pattern was markedly more diverse in WT animals while remaining flat call-dominated in *Fmr1* KO pups (Fig 4C,D). To quantify this difference, we next examined graph density, which measures the proportion of observed call-to-call transitions out of every theoretically possible combination (Fig. 4E). In both genotypes, graph density showed a clear developmental increase from p3 through p10, indicating a progressive expansion of syntactic flexibility during the peak period of USV production (Two-Way ANOVA (PND): F = 8.3554, **p = 0.0010). However, this developmental trajectory was significantly attenuated in *Fmr1* KO pups (Fig. 4E; Two-Way ANOVA (Genotype): F = 5.057, *p = 0.0306). These findings parallel observations in other ASD rodent models in which call type repertoire and sequencing are disrupted independently of acoustic properties^39,42^. Together, these results suggest that syntactic flexibility of vocal output may be a particularly sensitive measure of the neurodevelopmental consequences of *Fmr1* deletion during the early postnatal period.

## DISCUSSION

Deficits in communicative behavior are a hallmark of FXS, yet how FMRP loss shapes the developmental trajectory of vocal output has remained unclear. Here we characterized isolation-induced USVs in *Fmr1* KO and WT rat pups across postnatal days 3 to 21, providing a comprehensive developmental profile spanning acoustic properties, call output, temporal organization, and syntactic structure. The results reveal a clear dissociation: acoustic and temporal properties of individual calls are largely preserved in *Fmr1* KO pups, while the typical developmental increase in call number and expansion of vocal syntactic flexibility were significantly perturbed. These findings position syntactic organization, rather than physical acoustic properties, as the primary dimension of early vocal communication disrupted by loss of FMRP.

Our results indicate that the acoustic properties of isolation-induced USVs are dynamically regulated across development (Fig 2). However, we found no difference in call properties between WT and *Fmr1* KO littermate pups at any developmental timepoint. The preservation of acoustic call properties across the full developmental window is broadly consistent with the FXS human literature, in which phonological and articulatory abilities are relatively spared while higher-order aspects of communication are more severely affected^43^. These results appear to contrast with reports from *Fmr1* KO mouse models where spectral and temporal differences in pup USVs have been identified^29,44^, but this may reflect species differences in vocalization production rather than a genuine inconsistency in the effects of *Fmr1* deletion. Rats and mice differ substantially in the breadth and organization of their vocal repertoire, as well as in the anatomical substrates of vocal production^15,45^. These species-specific differences in vocal production machinery may confer differential sensitivity to FMRP loss at the level of individual call acoustics.

*Fmr1* KO pups exhibited reduced number of calls prior to ear opening relative to WT animals (Fig 3), in agreement with prior reports in Fmr1 KO mice^29,44^, and with the broader pattern of reduced communicative initiation in individuals with FXS^43^. Notably, the direction of this effect is the opposite to what has been reported in a *Tsc2*^+/-^ mouse model of Tuberous Sclerosis Complex (TSC), another common monogenic form of ASD, were elevated call rates are observed^46^. Intriguingly, it has been previously shown that TSC and FXS rodent models exhibit opposing changes to synaptic and cellular function^47^. Our current results suggest these divergent cellular changes may extend to communicative behavior. Crucially, the reduction in call quantity observed in *Fmr1* pups did not translate into broader impairment of temporal structure in calling behavior, with inter-call interval, bout duration, and sequences per bout preserved across all developmental timepoints. This result indicates that the mechanisms governing the rhythmic and structural patterning of distress signaling remain intact in *Fmr1* KO animals.

The most striking finding from this study is the altered developmental trajectory of vocal syntactic flexibility in *Fmr1* KO pups (Fig 4). In WT animals, the diversity of call-to-call transitions increased progressively from p3 through p10, reflecting a normal expansion of sequential behavioral repertoire during the peak calling period. This developmental expansion was significantly altered in *Fmr1* KO pups, whose transition networks remained comparatively static across the same window. This is consistent with the broader language phenotype in FXS individuals, which manifests primarily as delays in the development of pragmatic and sequential aspects of language: discourse coherence, topic maintenance, narrative structure, and flexible communicative use in context^43,48^. The parallel between attenuation of syntactic flexibility in rat pup USVs and impaired sequential communicative organization in ASD is further supported by evidence from other ASD rodent models in which call-to-call transition structure is disrupted independently of call number or acoustic properties^39,44,46^. Taken together, these findings identify vocal syntax as a sensitive and previously underappreciated readout of early communicative development in genetic models of ASD. Future work must determine whether interventions targeting this early postnatal window can restore the normal developmental trajectory of syntactic flexibility in FXS.

## MATERIALS AND METHODS

### Subjects

All experiments were approved by the Institutional Animal Care and Use Committee (IACUC) at the University of Illinois and carried out in compliance with guidelines and regulations specified by IACUC protocol #23223. Male wildtype (WT) rats (Charles River) were bred to heterozygous female *Fmr1* KO rats on an outbred Long-Evans background (LE-Fmr1em2Mcwi) to generate the male *Fmr1* KO and WT littermates used in this study. Founder *Fmr1* animals were obtained from the Medical College of Wisconsin and a colony was established and maintained in-house. Male pups were used exclusively, as FXS occurs more frequently and with greater severity in males due to the X-linked nature of the disorder^49^. Genotyping was performed by Transnetyx (Cordova, TN) from ear tissue collected at the time of weaning. All behavioral scoring and data analysis were performed with the experimenter blinded to genotype. Litters remained with the dam from birth through the completion of all recording sessions. All animals were housed in a colony room maintained at 22°C on a 12-hour light-dark cycle with ad libitum access to food and water.

Pups from a total of 4 litters were used for data collection across the developmental recording series, with sessions conducted at postnatal days 3, 6, 10, 14, and 21. Although the same litters contributed data at each timepoint, a strictly longitudinal design could not be maintained for all subjects due to early-stage sexing difficulty and logistical attrition. This semi-longitudinal approach nonetheless allowed for developmental trajectories to be characterized across the full p3–p21 window while minimizing cumulative disturbance to the home cage. The number of animals contributing data at each timepoint was as follows: p3 (7 WT, 9 KO), p6 (5 WT, 10 KO), p10 (10 WT, 6 KO), p14 (5 WT, 5 KO), and p21 (4 WT, 6 KO).

### Isolation Set-Up & Vocalization Recordings

At each developmental timepoint, rat pups were separated from the dam and placed into a holding cage within a double-walled sound isolation chamber. Ultrasonic vocalizations were then collected with an ultrasonic microphone (Avisoft UltraSoundGate CM16/CMPA) for 5-minutes, after which pups were then returned to the dam in the home cage. Call detection and acoustic feature extraction were performed using DeepSqueak (v.2.6.2), an automated USV detection and analysis platform based on a region-based convolutional neural network^33^.

## Data Analysis

### USV analysis pipeline

Initial call detection, acoustic properties, and call type classification were performed with DeepSqueak (v.2.6.2). All detected calls were first manually reviewed for artifact removal. Acoustic properties were then extracted directly from DeepSqueak’s built-in measurement pipeline applied to each call’s spectrogram. All further analyses were performed using a custom Python (3.10) pipeline. For each animal at each timepoint, calls were sorted chronologically by onset time within each isolation session. Inter-call intervals (ICIs) were computed as the time elapsed between the end time of call i and the begin time of call i+1 for all consecutive call pairs. Calls were then segmented into bouts and sequences using established temporal criteria^12,13^. A bout was defined as a continuous period of calling in which no two consecutive calls were separated by more than 2 seconds; an inter-call gap exceeding this threshold was taken to indicate the end of the current bout and the beginning of a new one. Within each bout, sequences were defined as runs of calls separated by no more than 500 ms; an inter-call gap exceeding this threshold within a bout was taken to indicate a new sequence. Per-animal metrics derived from this segmentation included total call number, mean ICI, mean bout duration, and mean number of sequences per bout.

### Call type classification

Call type labels were assigned using the DeepSqueak region-based convolutional neural network classifier, retrained on a held-out subset of recordings not used in the primary analysis. Ground truth labels for classifier training were generated by a trained scorer blinded to genotype and timepoint, who classified calls using a standardized decision tree based on established spectrotemporal criteria (Fig 4A)^11,12^. Classifier performance was validated against the hand- scored labels prior to application to the full dataset.

### Call types and syntax

For each animal at each timepoint, the proportion of calls belonging to each call type was computed as the number of calls of that type divided by the total number of calls in that session. To characterize the sequential structure, or syntax, of USV output, a directed transition probability matrix was constructed for each animal at each timepoint. For each consecutive call pair within a bout, a transition from the call type of call i to the call type of call i+1 was recorded. Raw transition counts were aggregated into a n × n matrix, where n is the number of call types, and each row was normalized by its sum to yield conditional transition probabilities, such that each entry P(i→j) represents the probability of transitioning to call type j given that the current call is of type i. Rows with zero total counts (call types not produced by that animal at that timepoint) were assigned uniform zero probability. A directed graph was then constructed from the resulting probability matrix for each animal and timepoint, with a directed edge included between call types i and j (excluding self-loops) wherever P(i→j) > 0. Graph density was computed as the number of observed edges divided by the total number of possible directed non-self-loop edges n(n − 1), as implemented in the NetworkX library (Hagberg et al., 2008), yielding a value between 0 and 1 representing the proportion of possible call type transitions that were utilized in that session.

### Statistical Analyses

Statistical analyses were performed in Python (3.10) using the scipy, statsmodels, and networkx libraries. For all USV metrics, the animal was treated as the unit of analysis; individual call measurements were averaged to the animal level prior to any statistical comparison to avoid pseudoreplication. Normality was assessed at each group-by-timepoint cell using the Shapiro- Wilk test applied both to the raw data and to the residuals of the fitted two-way model. Homogeneity of variance across groups was assessed using Levene’s test. Where normality and equal variance assumptions were satisfied, a two-way ANOVA with genotype and postnatal day as factors was used to assess main effects and their interaction, followed by Tukey HSD post-hoc comparisons where a significant main effect of genotype was detected. Where normality or equal variance assumptions were violated, the Scheirer-Ray-Hare test was used as a non-parametric two-way equivalent, decomposing ranked data into genotype, postnatal day, and interaction components with H statistics evaluated against a chi-squared distribution. When a significant main effect of genotype was identified, per-timepoint pairwise comparisons were conducted using Mann-Whitney U tests with Benjamini-Hochberg false discovery rate correction applied across timepoints; pairwise comparisons were not performed in the absence of a significant main effect. All values are reported as means ± SEM.

## DATA AVAILABILITY

Analysis code and pre-processed data available at Zenodo (https://doi.org/10.5281/zenodo.20940042). Raw data available upon reasonable request.

